# Synergistic Effects of Sulopenem in Combination with Cefuroxime or Durlobactam against *Mycobacterium abscessus*

**DOI:** 10.1101/2023.12.15.571879

**Authors:** Khalid M. Dousa, Eunjeong Shin, Sebastian G. Kurz, Mark Plummer, Mary Nantongo, Christopher R. Bethel, Magdalena A. Taracila, David C. Nguyen, Barry N. Kreiswirth, Charles L. Daley, Kenneth E. Remy, Steven M. Holland, Robert A. Bonomo

## Abstract

*Mycobacterium abscessus* (*Mab*) affects patients with immunosuppression, Cystic Fibrosis (CF), or underlying structural lung diseases. Additionally, *Mab* poses clinical challenges due to its resistance to multiple antibiotics. Herein, we investigated the synergistic effect of dual β-lactams [sulopenem and cefuroxime (CXM)] or the combination of sulopenem and CXM with a β-lactamase inhibitors [BLI; avibactam (AVI) or durlobactam (DUR)]. The sulopenem-CXM combination yielded low minimum inhibitory concentration MIC values for 54 clinical *Mab* isolates and ATCC19977 (MIC_50_ and MIC_90_ ≤ 0.25 μg/mL). Similar synergistic effects were observed in time-kill studies conducted at concentrations achievable in clinical settings. Sulopenem-CXM outperformed monotherapy, yielding ∼1.5 Log_10_ CFU/mL reduction during 10 days. Addition of BLIs enhanced this antibacterial effect, resulting in additional reduction of CFUs (∼3 Log_10_ for sulopenem-CXM and AVI and ∼4 Log_10_ for sulopenem-DUR). Exploration of the potential mechanisms of the synergy focused on their interactions with L,D-transpeptidases (LDTs; LDT_Mab1_–LDT_Mab4_), Penicillin-Binding-Protein B (PBP-B), and D,D-Carboxypeptidase (DDC). Acyl complexes identified via mass spectrometry analysis, demonstrated the binding of sulopenem with Ldt_Mab2_-Ldt_Mab4_, DDC, and PBP B, and CXM with Ldt_Mab2_ and PBP-B. Molecular docking suggested formation of a covalent adduct between sulopenem and Ldt_Mab2_ after the nucleophilic attack of the cysteine residue at the β-lactam carbonyl carbon, leading to the cleavage of the β-lactam ring, and the establishment of a thioester bond linking the Ldt_Mab2_ with sulopenem. In conclusion, we demonstrated the biochemical basis of the synergy of sulopenem-CXM with or without BLI. These findings potentially broaden selection of oral therapeutic agents to combat *Mab*.

## INTRODUCTION

*Mycobacterium abscessus* (*Mab*), a non-tuberculous mycobacterium (NTM), is well-known for its recalcitrance to treatment, presenting a formidable challenge for clinicians (1). Eradication of *Mab* infection is challenging given its innate resistance to most anti-tuberculous medications, proclivity to form biofilms, need for multi-drug regimen with associated toxicity, longer duration of therapy, predilection for infecting immunocompromised patients with underlying lung diseases, and the challenge to mount an effective immune response to clear the infection (2). In the presence of macrolide resistance genes (*erm41* and *rrl*) in subsp. *abscessus*, induced or acquired respectively, treatment failure rates can soar up to 70% in some reports, exceeding those observed with multidrug-resistant tuberculosis (MDR-TB) (3, 4). In light of these challenges, significant research efforts have focused on identifying essential genes and targets for drug development, including the repurposing of β-lactams alone or in combination therapies.

*Mab* contains a class A β-lactamase enzyme, known as Bla_Mab_ (5), which is capable of hydrolyzing penicillins, and cephalosporins. Penems appear to be more resistant to hydrolysis (5). Despite the presence of Bla_Mab_, current treatment guidelines do not address this issue (6). Older β-lactamase inhibitors (BLIs), such as clavulanic acid and sulbactam, are ineffective against Bla_Mab_ (5). However, recent research has identified that the newly developed diazabicyclooctane (DBO) class of BLIs (avibactam; AVI, nacubactam, and zidebactam) exhibits activity against Bla_Mab_, with some demonstrating intrinsic antimicrobial activity whose clinical significance is yet to be determined (7–12). Durlobactam (DUR) is a DBO compound with expanded spectrum of activity compared to other DBO’s, demonstrating robust inhibition against class A, C, and D serine β-lactamases. DUR’s potential to enhance *in vitro* susceptibility when combined with β-lactams, accompanied by the elucidation of a biochemical rationale underlying its mode of action was previously assessed (7).

The peptidoglycan structure of *Mab* is notably divergent from that of most bacterial species, primarily owing to its synthesis involving both L,D-transpeptidases and penicillin-binding proteins (PBPs), which include two protein classes: D,D-transpeptidases and D,D-carboxypeptidase. D,D-transpeptidases play a pivotal role in the synthesis of peptidoglycan, while D,D-carboxypeptidases are responsible for catalyzing the removal of terminal amino acids from peptidoglycan sidechains. Several essential PBPs, including PBP-lipo (MAB_3167c), PBP B (MAB_2000), DacB1, and DDC, were investigated, revealing that their “knockout” resulted in a synergistic growth inhibition effect (8–10).

Most recently, the role of combination β-lactams in inhibiting L,D-transpeptidation, the major peptidoglycan crosslink reaction in *Mab* has become an area of intense investigation. The five L,D-transpeptidases (LDT_Mab1-5_) (11), which are essential enzymes for catalyzing transpeptidation during cell wall synthesis, were previously studied and was found to be inhibited by carbapenems and cephalosporins but not penicillins (12). These reports suggested that by combining two β-lactams with or without a β-lactamase inhibitor, multiple targets in cell wall synthesis pathway can be inactivated (7, 11, 13), leaving the cell wall in a compromised state. The synergistic effects of dual β-lactams (imipenem and ceftaroline combination) (11), was achieved through the inhibition of multiple target proteins. Emerging clinical data further corroborate the effectiveness of dual β-lactam combinations (14–16). However, the co-administration of imipenem and cefoxitin, which are the sole two β-lactams recommended in the guidelines for *Mab* treatment, has undergone clinical trials with outcomes that lack clear evidence of efficacy (17). These observations indicate the necessity for an optimized β-lactam treatment strategy and lend support to the hypothesis that achieving optimal β-lactam therapy may entail the simultaneous targeting of multiple proteins within the intricate and highly redundant network of enzymes participating in peptidoglycan biosynthesis.

Sulopenem, formerly known as CP-70,429, is a novel broad spectrum thiopenem β-lactam compound with a structure similar to other carbapenems, but with modifications that provide enhanced stability and activity against several β-lactamase enzymes (18). The molecule has small molecular weight (345 Da), contains a β-lactam ring with a hydroxyethyl group attached at C6 and a thiazolidine ring fused to it (**FIGURE 1**). The side chain attached to the C2 possess a sulfoxide group which enhances the compound’s activity against Gram negative bacteria such as the *Enterobacterales* (18, 19). These structural features are unique and make it a potential new treatment option for drug-resistant bacterial infections. Sulopenem safety and efficacy are currently being evaluated in phase 3 clinical trial (NCT03357614, entitled "Sulopenem for the Treatment of Complicated Urinary Tract Infections (SUNSHINE)). In 2020, sulopenem received Qualified Infectious Disease Product (QIDP) designation from the US FDA. Due to its availability in oral formulation, Sulopenem offers clear advantages for the patient and for compliance with the dosing regimen, however, it necessitates co-administration with probenecid to mitigate renal clearance, as its oral bioavailability typically falls within the range of 30% to 40% (18).

**FIGURE 1:**
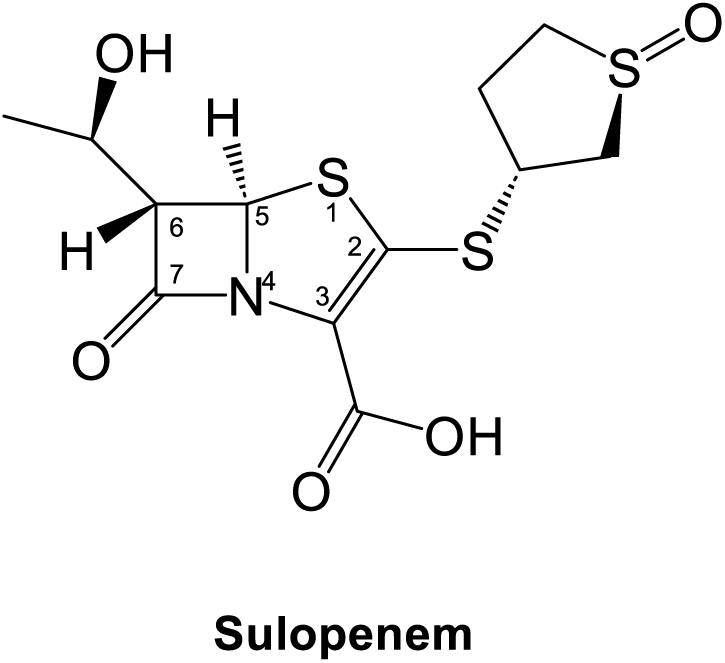
Sulopenem chemical representation

In this study, we used cell-based, and *in-vitro* static concentration time kill (SCTK) assay and other biochemical techniques to extend the previous investigation that examined imipenem and ceftaroline. We assessed the synergistic activity of sulopenem and cefuroxime and we also tested whether the presence or absence of β-lactamase inhibition would further demonstrate improved susceptibility. Our hypothesis is that sulopenem, an oral carbapenem antibiotic targets multiple L,D-transpeptidases, Ldt_Mab2_ -Ldt_Mab4_, DDC, and PBP B. The conserved active site histidine activates the catalytic site cystine to drive acyl enzyme formation with sulopenem. This results in significant improvement microbial killing of *Mab* both in the presence or absence of β-lactamase inhibition.

## RESULTS

### Sulopenem exhibits *in vitro* activity comparable to imipenem

An initial evaluation of the *in vitro* antimicrobial activity of sulopenem was conducted on the *Mab* ATCC 19977 strain, using Middlebrook 7H9 broth supplemented with 10% (vol/vol) oleic albumin dextrose catalase and 0.05% (vol/vol) Tween 80. The minimal inhibitory concentration (MIC) for sulopenem alone was found to be 1 µg/ml. Further susceptibility testing with sulopenem against a set of 54 previously characterized *Mab* subsp. *abscessus* clinical isolates demonstrated MICs similar to the ATCC 19977, with MIC_50_ of 2 µg/ml and MIC_90_ of 4 µg/ml (**Table 1**). These values are comparable to the previously reported MIC_50_ and MIC_90_ of imipenem (11).

**TABLE 1:** Comparative MIC_50_s and MIC_90_s for 54 *Mab* Clinical Strains and ATTCC 19977: Evaluating Sulopenem (SUL), Cefuroxime (CXM), and Combined Sulopenem with 4 µg/ml Cefuroxime.

| Antibiotics (µg/ml) | MIC range (µg/ml) | MIC<sub>50</sub> (µg/ml) | MIC<sub>90</sub> (µg/ml) |
| --- | --- | --- | --- |
| CXM | 8-64 | 16 | 32 |
| SUL | 1-8 | 2 | 4 |
| SUL + CXM (4 µg/ml) | ≤ 0.25 - 8 | ≤ 0.25 | ≤ 0.25 |

### Checkerboard assay reveals 4 µg/ml CXM concentration as synergistic in combination with sulopenem

A checkerboard assay was conducted to assess the potential synergistic effect of sulopenem and CXM against *Mab* strain ATCC 19977. The checkerboard assay was designed to simultaneously evaluate the inhibitory activities of both sulopenem and CXM in a two-dimensional matrix, where serial dilutions of sulopenem were combined with CXM. The MIC of CXM against *Mab* ATCC 19977 was determined to be in 16-32 µg/mL range. Based on this MIC value, a concentration of 4 µg/mL CXM was selected for inclusion in the assay to evaluate its potential synergy with sulopenem.

### Synergistic effects of sulopenem and CXM combination against *Mab* subsp. *abscessus* clinical isolates

CXM susceptibility against the 54 clinical isolates was subsequently tested. As anticipated, CXM demonstrated diminished activity against *Mab* clinical isolates in comparison to sulopenem, with an MIC range of 8-64 µg/mL. The MIC_50_ and MIC_90_ values were 16 µg/mL and 32 µg/mL, respectively. The addition of CXM at a fixed concentration of 4 µg/mL significantly improved the potency of sulopenem by lowering the MICs. In fact, 52 of 54 isolates demonstrated an MIC of sulopenem less than 0.25 µg/mL in the presence of CXM, indicating a pronounced synergistic effect between the two antimicrobial agents (**FIGURE 2**).

**FIGURE 2:**
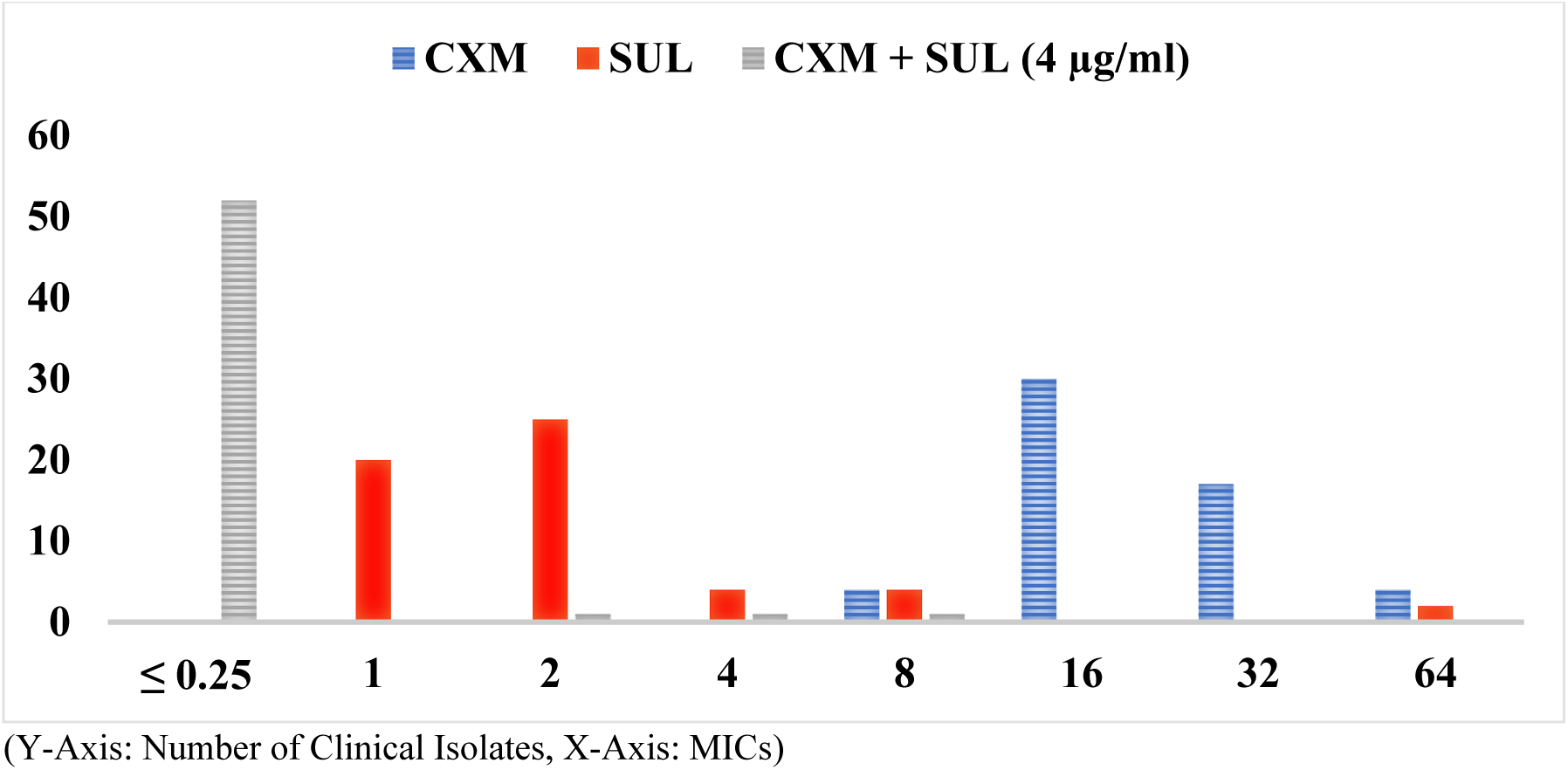
Minimal Inhibitory Concentration Distribution Comparison for sulopenem (SUL), Cefuroxime (CXM), and Combined Sulopenem with 4 μg/ml Cefuroxime Against 54 Mab Clinical Strains and the ATTCC 19977 Strain (See Table 1 for MIC Data)

### Synergistic effect of sulopenem in combination with CXM or BLIs in time-kill study

Bactericidal activity was assessed through the static concentration time-kill experiment. At 4 times the MIC (4 x MIC of sulopenem and CXM), sulopenem (8 µg/mL) and CXM (64 µg/mL) exhibited a 2-log reduction in Colony-Forming Units (CFU) after 2-3 days against ATCC 19977 (FIGURE 3). However, at 2 µg/mL of sulopenem and 8-12 µg/mL of CXM, both agents only achieved growth inhibition (stasis) without significant bactericidal effect. When combined, sulopenem and CXM demonstrated a slight synergistic effect, resulting in a 1-2 log CFU difference compared to monotherapy. Combining a fixed concentration of 8 µg/mL CXM with sulopenem or a fixed concentration of 2 µg/mL of sulopenem with CXM led to a reduction in bacterial load comparable to monotherapy. Nevertheless, during the course of 10 days, regrowth of bacteria was observed in most concentration ranges for both monotherapy and combination treatments.

**FIGURE 3:**
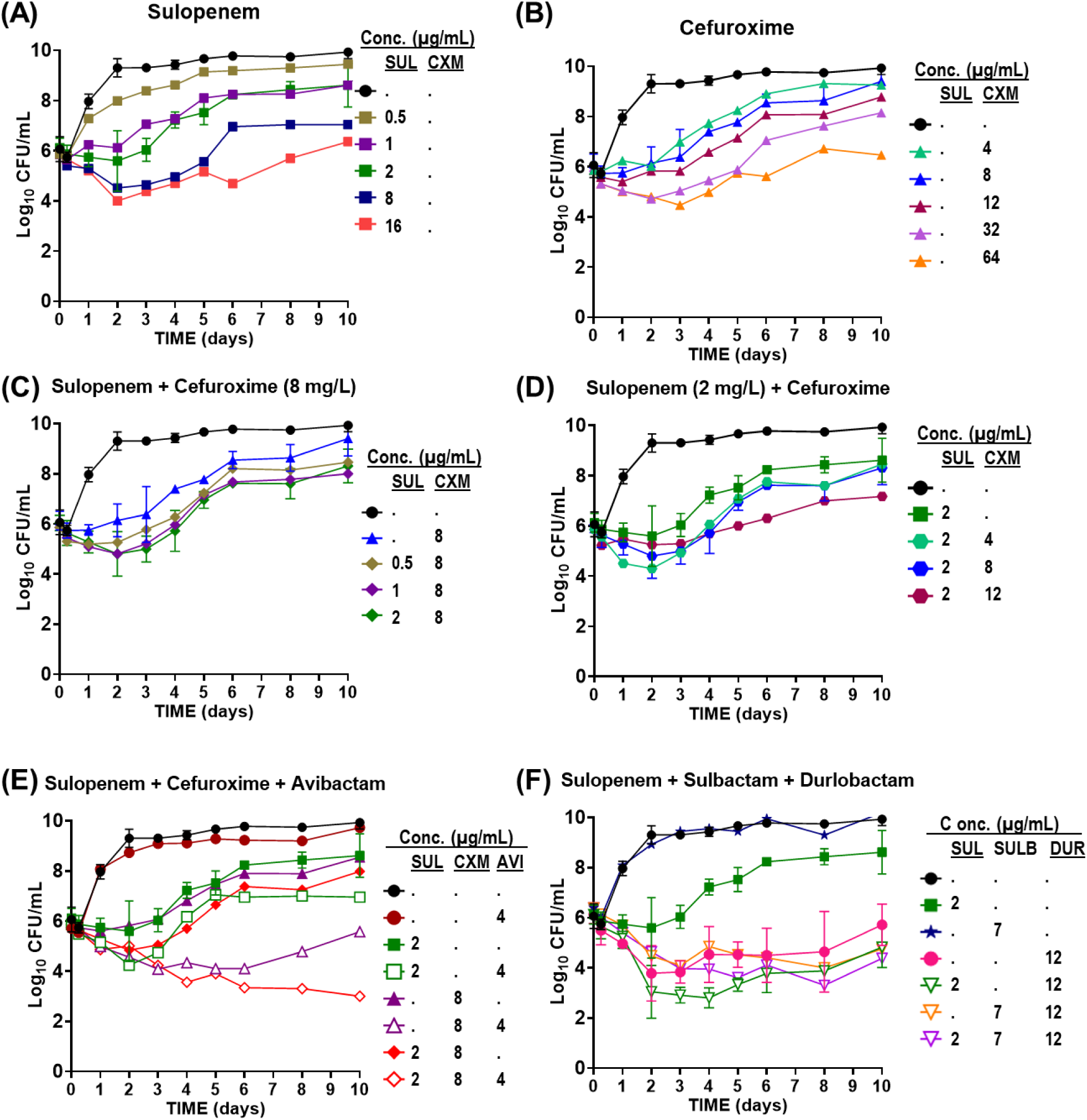
Time-kill curves of sulopenem (A) in monotherapy, cefuroxime (CXM) in monotherapy (B), the combination of sulopenem and CXM (C and D), sulopenem in the presence of avibactam (AVI) (E), and the combination of sulopenem and sulbactam with or without DUR (F) against ATCC 19977.

Addition of BLIs, AVI (4 µg/mL) or DUR (12 µg/mL), enhanced the bactericidal activity of sulopenem. The combination of sulopenem + AVI resulted in a decrease of up to 2-log CFU, while sulopenem + DUR exhibited a 3.5-log CFU reduction. Although 4 µg/mL of AVI alone did not lead to significant killing, the combination of AVI + sulopenem yielded a 2.7-log reduction in Log_10_ CFU unit. At 12 µg/mL (6 folds of MIC of DUR), DUR monotherapy achieved a 2-log reduction in bacterial load, suggesting possible intrinsic antimicrobial activity in addition to its BLI activity against *Mab*, which was more effective compared to AVI. While sulopenem +AVI showed regrowth on day 4, Sulopenem + DUR inhibited regrowth successfully for up to 10 days.

Considering that DUR is commercially available with sulbactam, we also investigated the bacterial effect of triple combination of sulopenem + sulbactam + DUR. Sulbactam had no bactericidal effect either alone or in combination with sulopenem or DUR. However, when combined with sulopenem and DUR, sulbactam exhibited a similar synergistic effect on bacterial killing. Furthermore, SCTK studies were conducted using two randomly selected clinical isolates (122 and 686). Comparable results were obtained for isolate 122 (**Supplement Fig. 2**), whereas less killing was observed for isolate 686 (**Supplement Fig. 3**).

### Timed electrospray ionization mass spectrometry (ESI/MS) captured covalent adduct formation between Ldt_Mab2-4_, D,D-Carboxypeptidase, PBP B, and Bla_Mab_ with sulopenem and cefuroxime

To explore potential mechanism for transpeptidation inhibition by sulopenem, we investigated if sulopenem and CXM could form acyl complex with Ldt_Mab1_ – Ldt_Mab4_, D,D-carboxypeptidase, and PBP B. Ten micrograms of Ldts, D,D-carboxypeptidase, and PBP B were incubated with sulopenem and CXM at room temperature in a molar ratio of 1:20 (enzyme to sulopenem) for 5 min, 2 h, and 24 h in 50 mM Tris-HCl (pH 7.5) and 300 mM sodium chloride for a total reaction volume of 20 μl. Sulopenem underwent reaction with Ldt_Mab2_, Ldt_Mab3_, Ldt_Mab4_, D,D-carboxypeptidase (**FIGURE 4, 7 and Supplement Fig. 1**) and PBP B The resultant covalent drug adduct was measured using intact-protein ultra-performance liquid chromatography (UPLC) coupled with EI/MS analysis. A peak corresponding to the mass of Ldt_Mab2_ or Ldt_Mab4_ plus +86 Da mass shift was captured. This mass difference corresponds to the addition of the sulopenem followed by post-acylation changes and fragmentation of the compound. This phenomenon stems from the distinctive cleavage of the C5-C6 bond, a mechanism previously reported in penems and carbapenems (20–23). Sulopenem formed a +349 Da adduct with Ldt_Mab3_, D,D-carboxypeptidase, and PBP B. In contrast to sulopenem, CXM engaged solely with Ldt_Mab2_ and PBP B, resulting in the formation of a +381 Da adduct for Ldt_Mab2_ and a +364 Da adduct for PBP B (**FIGURE 4**).

**FIGURE 4:**
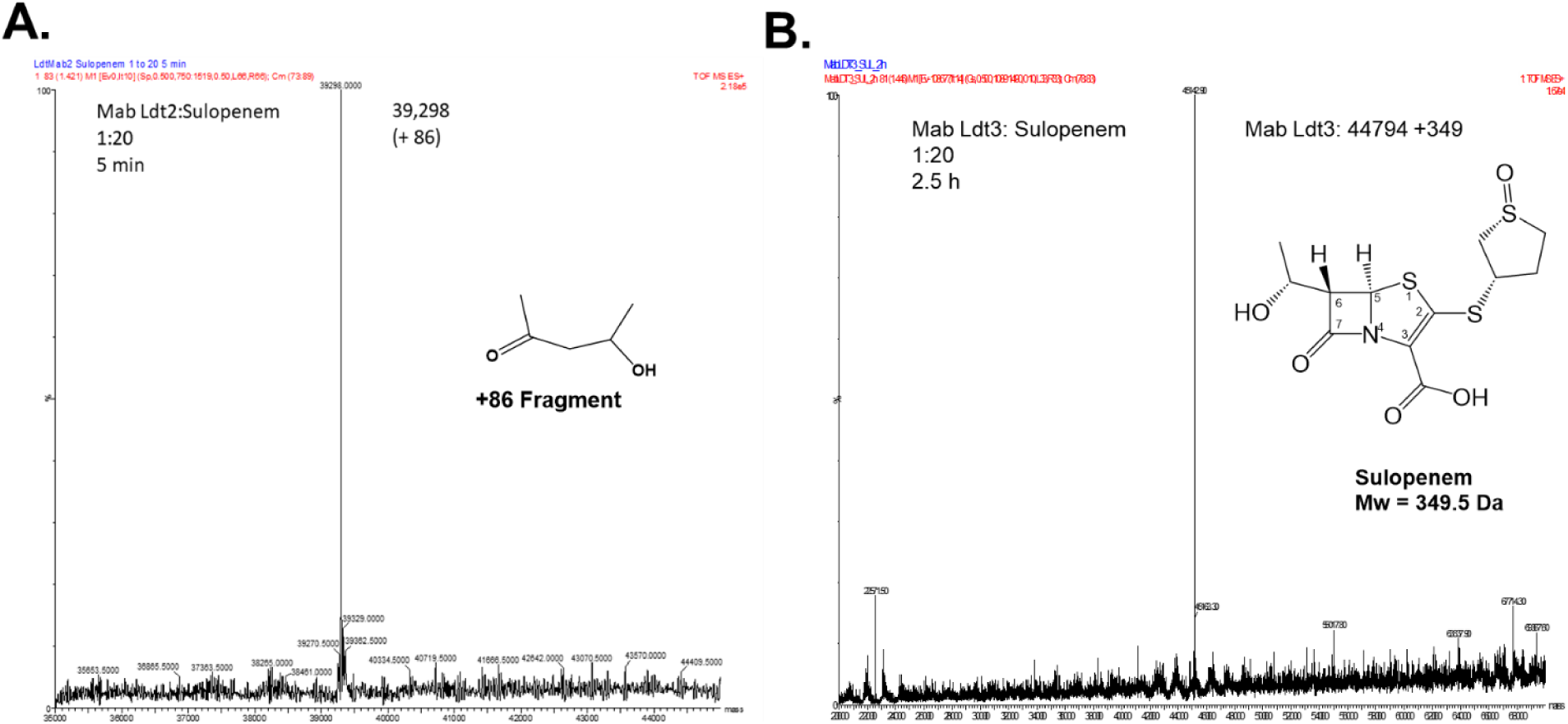
Capturing covalent adduct formation between sulopenem and Ldt_Mab2_ (A) and Ldt_Mab3_ (B) using Timed Electrospray Ionization Mass Spectrometry. After 5 min incubation of sulopenem with Ldt_Mab2_ the adduct formed is 86 Da (A). When incubated with LdtMab_3_ a 349 Da (sulopenem Mw) adduct is preserved after 2.5h (B).

We subsequently examined the covalent binding of sulopenem and CXM to Bla_Mab_. Following co-incubation for time intervals of 5 seconds, 15 seconds, and 2 hours, we were unable to detect any interaction between sulopenem and/or CXM with Bla_Mab_. At these experimental conditions, Bla_Mab_ (28,433.5 Da) demonstrated binding with AVI (+265.5 Da) and DUR (+277 Da), as previously reported (7, 11).

### Computational modeling and molecular docking of sulopenem with Ldt_Mab2_ and Ldt_Mab3_

For the molecular docking of sulopenem, 2 different models of Ldt_Mab2_ enzyme were used (11): *i*) closed cavity, observed in apo enzyme structure; and *ii*) the model with the opened active site, observed in crystal structures with the compounds trapped into the active site.

The automatic DOCKER module did not generate any conformations with the sulopenem intact. The Michaelis-Menten complex generated when the open active site Ldt_Mab2_ conformation was chosen **(FIGURES 5A, C)** show His333 positioned very close to the catalytic Cys351: HG, (His333:NE2 less than 2.5Å). The imidazole ring of His333 is held into this productive conformation through a network of hydrophobic interactions by Val319 and Ala335. Once sulopenem is docked in the active site, the His333 imidazole ring is further stabilized by the hydroxyethyl moiety which makes interactions with Trp337. This proximity is consistent with previous studies (11), which show that the His333:NE2 is necessary to activate Cys351 for the nucleophilic attack on the lactam bond of sulopenem (**SCHEME 1**). The carbonyl group of sulopenem is positioned into the oxyanion hole created by Cys351:NH and Gly350:NH (**FIGURE 5A**). After the acyl formation complex, the His333 imidazole ring moves outside (≈ 5.6-6Å) from the oxyanion hole **(FIGURE 5B)**. The generated molecular docking complex of the acyl sulopenem and Ldt_Mab2_ show the sulopenem carbonyl positioned outside of the oxyanion hole (**FIGURE 5B and C**).

**SCHEME 1:**
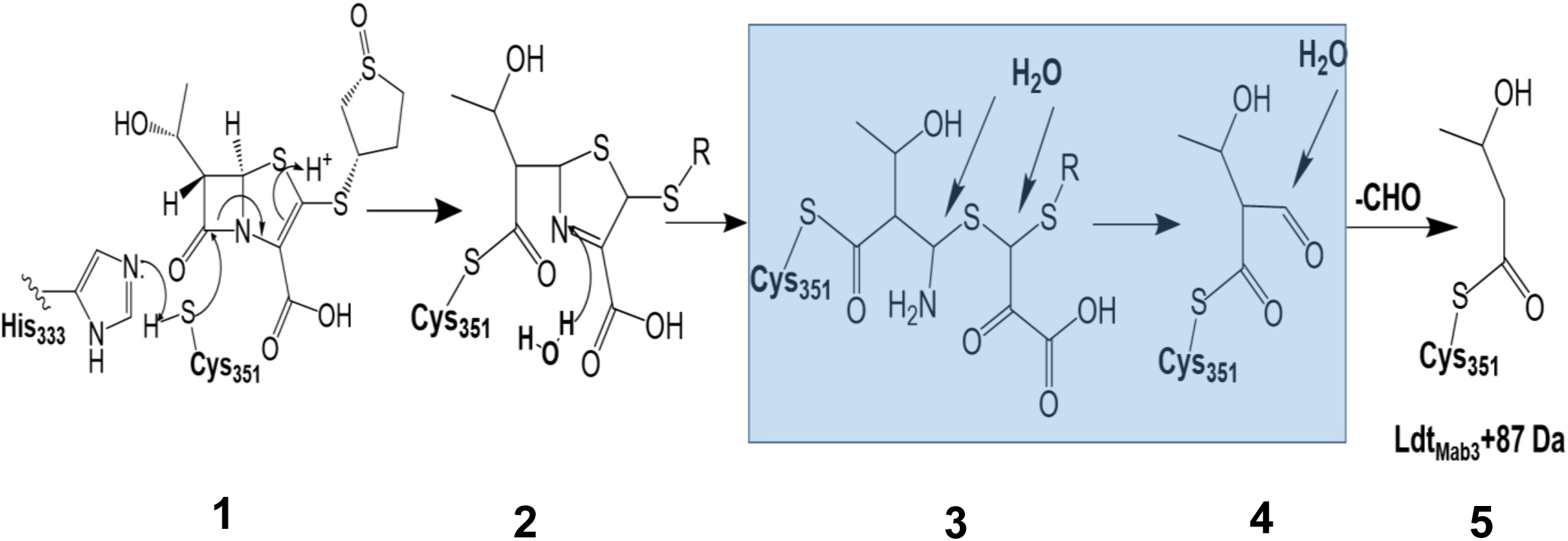
Proposed mechanism of action between sulopenem and Ldt_Mab_ transpeptidases. The covalent adduct formation between sulopenem and Ldt_Mab2_ can be explained through the nucleophilic attack of the cysteine residue at the carbonyl carbon of the β-lactam ring in sulopenem, with the help for activation from His333 (1). This nucleophilic attack results in the opening of the β-lactam ring and the formation of a thioester bond between the enzyme and sulopenem (**2**). Steps (3) and (4) are intermediate steps in the reaction mechanism suggested by the MDS results, where 3 water molecules were recruited into the active site. The end adduct of 87 Da (5) was observed after 5min incubation of Ldt_Mab2_ with sulopenem. When sulopenem is incubated with Ldt_Mab3,_ after 2.5h the adduct with the intact sulopenem (2) was observed in the MS. The molecular modeling of sulopenem and Ldt_Mab3_, does not result in water molecules recruited into the active site. This may suggest that the reaction mechanism of sulopenem with Ldt_Mab3_ ends after step 2, with the release of the intact sulopenem.

**FIGURE 5:**
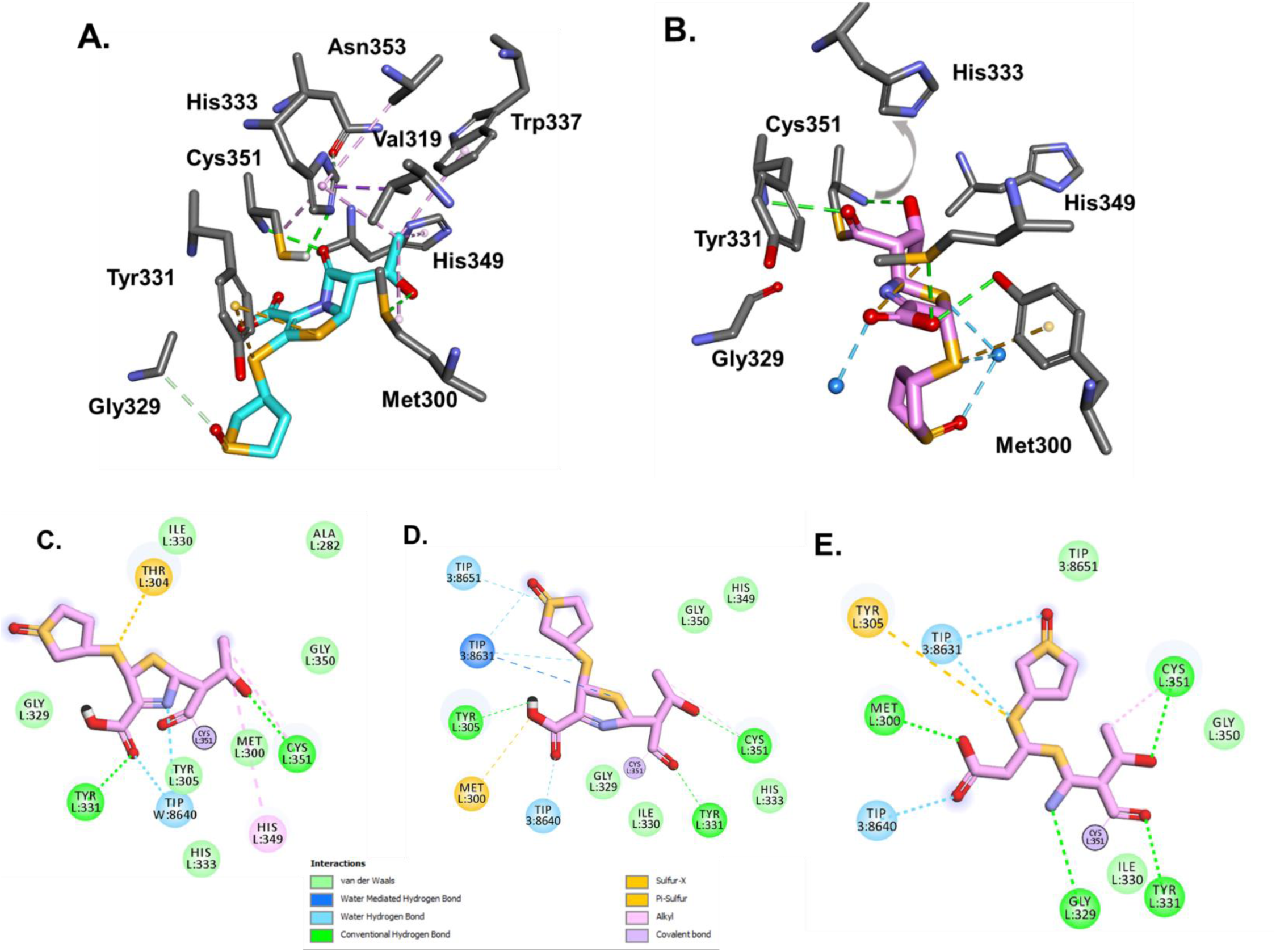
Ldt_Mab2_ and sulopenem as Michaelis-Menten (A) and acyl enzyme (B) complexes. During MDS simulation of the acyl-enzyme (C, D) and sulopenem fragment (E) complexes, three water molecules are recruited into the active site of Ldt_Mab2_. Initially, (A) H333 is at H-bond distance from Cys351and ready to activate it for acyl-enzyme formation. After the acyl formation (B), the His333 is moving away from Cys351, and sulopenem carbonyl is positioned outside of the oxyanion hole. The potential H-bonds interactions are represented with green, hydrophobic interaction with pink, and sulfur with yellow.

To further understand the potential interactions of sulopenem and explore/explain the adduct formation of +87 Da seen on the ES/MS data **(FIGURE 4)** when Ldt_Mab2_ is incubated for 5min with sulopenem, the molecular dynamic simulation was run for the acyl complex **(FIGURE 5C, D)** and potential fragment of sulopenem **(FIGURE 5E)**. The model shows three water molecules recruited into the active site during the MDS. The first water molecule is positioned H-bond distance from thiazolidine ring N4, **(SCHEME 1 and FIGURE 5C)** making possible interaction with moieties on the sulopenem fragment in the active site. Two more water molecule that are recruited are less than 3Å from sulfa groups from the acyl and/or fragmented sulopenem **(FIGURE 5D and E)**.

The mass spectrometry data shows that Ldt_Mab3_ forms an intact adduct with sulopenem, after 3 h incubation **(FIGURE 4B)**. The molecular modeling and the docking of sulopenem into the active site of Ldt_Mab3_ show that the carbonyl is positioned toward the oxyanion hole formed by Cys351:NH and Gly350:NH, ready for acylation. However, His333:NE2 is more than 5Å away from the catalytic Cys351 **(Figure 6A and B)**, and the sulopenem carbonyl makes H-bonds with Gly329. This suggest that the acyl enzyme complex formation may take longer to form (slow acylation). When the acyl enzyme is formed, the complex is held and stabilized in the active site of Ldt_Mab3_ through a network of H-bonds and hydrophobic interactions **(FIGURE 6C, D and SCHEME 1)**.

**FIGURE 6:**
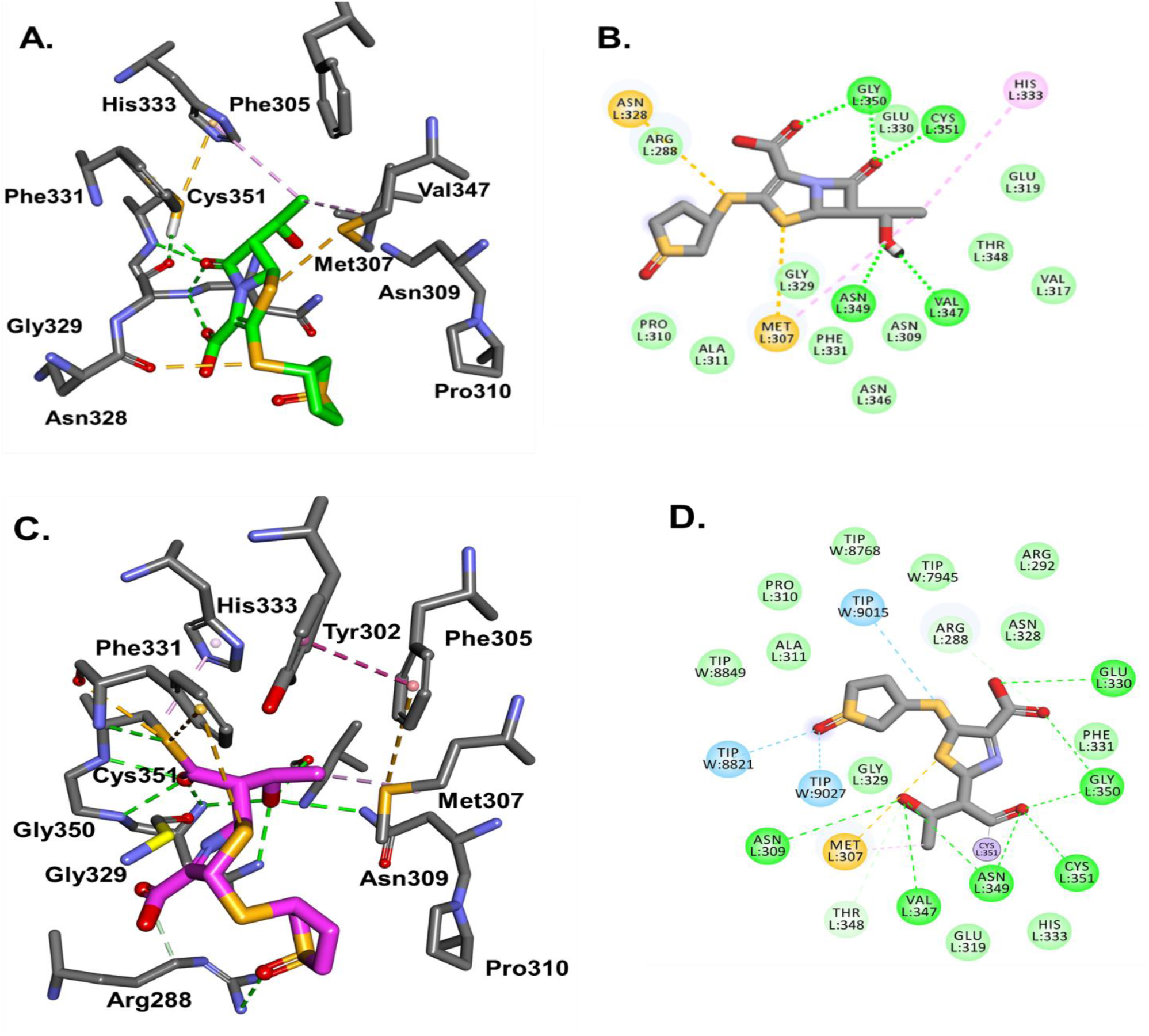
Ldt_Mab3_ and sulopenem molecular docking as Michaelis-Menten (A, B) and acyl enzyme (C, D) complexes. The sulopenem carbonyl is positioned toward the Ldt_Mab3_ oxyanion hole formed by Cys351:NH and Gly350:NH, ready for acylation. However, His333:NE2 is more than 5Å away from the catalytic Cys351, and the sulopenem carbonyl makes H-bonds with Gly329 (A, B). This suggest that the acyl enzyme complex may take longer to form. When the acyl enzyme is formed (C), the complex is held and stabilized in the active site of Ldt_Mab3_ through a network of H-bonds (green) and hydrophobic interactions (pink) (C, D).

This unique behavior of sulopenem can be explained due to the variability in the active site of Ldt_Mab2_ and Ldt_Mab3_ **(Supplement Fig.4)**. The variability is observed in the sequence alignment (residues variability **Supplement Fig.4F)**, and the size and shape of the “outside” and” inside” active site cavities **(Supplement Fig.4A, B).** The entrance of the active site cavity is similar for both enzymes, with an approximatively 12-13Å opening (**Supplement Fig.4A, B)**. However, the inside cavity increases from 5-6 Å in Ldt_Mab2_ to up to 14 Å in Ldt_Mab3_ **(Supplement Fig. 4A and B)**, mostly due to the changes from Tyr to Ala, and Tyr331 to Phe. The most important changes is observed at the entrance on the active site, with the Tyr305 in Ldt_Mab2_ replaced by alanine and proline 310, and His339 replaced by Asn349 **(Supplement Fig. 4)**. The hairpin loops in Ldt_Mab3_ have a 7a.a deletion **(Supplement Fig.4 A, B (yellow representation) and F)**. This deletion and the Pro310 in the middle of the alpha helix, changes the architecture of the active site dramatically, (**Supplement Fig.4C, D)** making it smaller and more restrained in Ldt_Mab3_. The Ldt_Mab3_ is closer in overall topology to Ldt_Mt5_ (24) and also found it to be structurally and functionally distinct.

## DISCUSSION

In recent years, the repurposing and reintroduction of β-lactams as a potential treatment for *Mab* infections has gained renewed interest. This interest has been augmented by the introduction of new DBO β-lactamase inhibitors. DBOs, such as AVI, relebactam, and the recently FDA-approved DUR have demonstrated the ability to inhibit the enzymatic activity of Bla_Mab_ and restore *in vitro* susceptibility to β-lactams in *Mab* (7, 25–27). For example, the addition of AVI was found to increase *Mab* susceptibility to ceftaroline, with even further reductions in MICs observed upon the addition of ceftazidime (26). Similarly, DUR enhanced the susceptibilities for CXM and imipenem in a large collection of clinical isolates of *Mab* subspecies *abscessus* (7). Relebactam was also found to enhance susceptibility when combined with amoxicillin and imipenem against *Mab* isolates (25).

The precise dynamics of these DBOs in Bla_Mab_ inhibition and inhibition of Ldts is complex. In fact, previous studies have demonstrated that dual β-lactam combinations can significantly reduce MICs, without the need for Bla_Mab_ enzyme inhibition. In time kill profiles of sulopenem in presence of AVI (**Supplement Fig. 2**), we found 1-1.5 Log_10_ reduction in CFU unit compared to absence of DBOs, while AVI did not show any killing against ATCC 19977 and clinical isolates *Mab*122. This suggested that the synergistic effect has been driven by their activity to inhibit Bla_Mab_. However, DUR yielded significant killing showing 2 log_10_ decrease of bacterial load by itself and 3 log_10_ reduction by combination with sulopenem, suggesting DUR enhanced *in vitro* bacterial killing by both inhibition of Bla_Mab_ and transpeptidation. Combination β-lactams was also found highly active *in vitro* in prior studies, this is exemplified by various combinations such as ceftaroline and imipenem (11), doripenem and cefdinir (12), ceftazidime and ceftaroline (28), imipenem and cefdinir, imipenem and cefoxitin (26), and sulopenem with CXM (29, 30), tebipenem, and CXM combined with amoxicillin (30) questioning the need for Bla_Mab_ inhibition. Clinicians have utilized this approach in treating macrolide resistant *Mab* infections with reports of clinical success (14–16).

The central question that persists is whether the additive or synergistic impact of DBOs on *Mab* susceptibility arises from their ability to inhibit Bla_Mab_, their inherent antimicrobial properties in hindering L,D-transpeptidases, or a combination of both factors? In our mass spectrometry data, we did not observe any evidence of binding between sulopenem and CXM with Bla_Mab_ (**FIGURE 7**). This observation aligns with previous reports indicating that the kinetic parameters for the interaction between CXM and Bla_Mab_ are exceptionally high, making it challenging to determine, thus suggesting that Bla_Mab_ may not hydrolyze CXM (5). Moreover, it is noteworthy that AVI and DUR exhibit inhibitory effects not only on Bla_Mab_ but also on LDTs (7). This multifaceted inhibitory activity may elucidate the observed synergistic effects of β-lactam antibiotics combined with β-lactamases, which stem from the additional role of DBOs in inhibiting LDTs. This is one possible explanation and further research will be necessary to elucidate the question.

**FIGURE 7:**
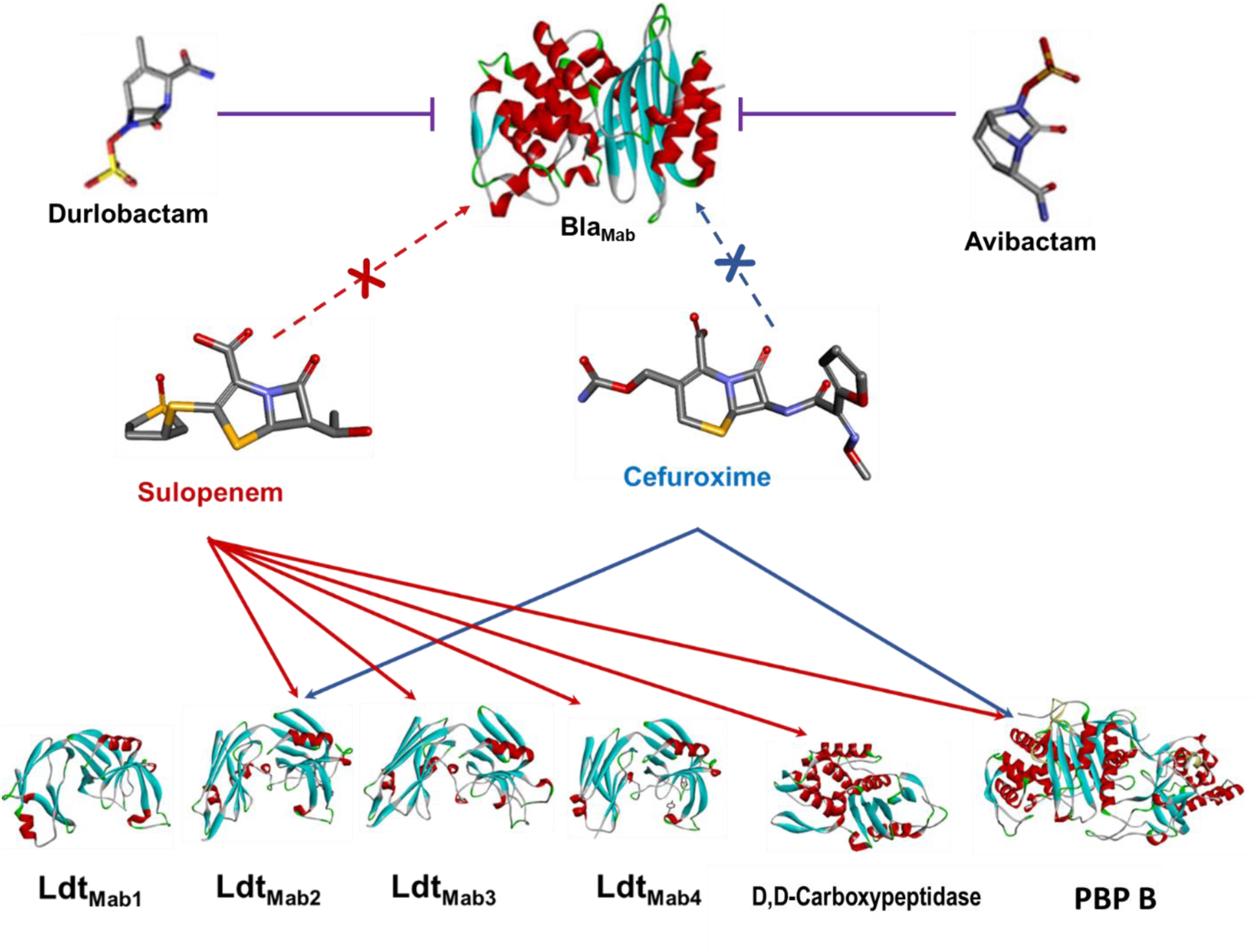
Interaction between Bla_Mab_, L,D-Transpeptidases (Ldt_Mab1_ to Ldt_Mab4_), D,D-carboxypeptidase, and PBP B with β-lactams (sulopenem and cefuroxime) and β-lactamase inhibitors (durlobactam and avibactam).

Ldt_Mab2_ belongs to a family of enzymes that catalyze the formation of 3→3 cross-links between peptidoglycan layers in the cell wall (20). In *Mab*, Ldt_Mab2_ plays a pivotal role in maintaining cell wall integrity and is essential for bacterial survival. The active site of Ldt_Mab2_ contains a highly conserved cysteine residue, which acts as a nucleophile during the catalytic process (11). The covalent adduct formation between sulopenem and Ldt_Mab2_ can be explained through the nucleophilic attack of the cysteine residue at the carbonyl carbon of the β-lactam ring in sulopenem (**SCHEME 1** and **FIGURE 5A)**. This nucleophilic attack results in the opening of the β-lactam ring and the formation of a thioester bond between the enzyme and sulopenem (**SCHEME 1** and **FIGURE 5B**).

Similarly, previous studies have demonstrated that tebipenem, another oral carbapenem, exhibits binding and activity against Ldt_Mab2_ (12). Like sulopenem, tebipenem is a penem antibiotic with a similar bicyclic ring system composed of a β-lactam ring fused to a five-membered ring, which is connected to an azetidine ring at C2. Based on the tebipenem molecular modeling and its binding to Ldt_Mab2_, the value of theoretical free energy of binding for Ldt_Mab2_ and tebipenem was higher when compared with doripenem free energy of binding, suggesting that tebipenem is a better inhibitor. (12).

The range of MIC values of sulopenem against clinical isolates in this cohort, which ranged from 1-8 μg/ml, is quite intriguing despite the absence of CLSI breakpoints for sulopenem activity against *Mab*. Currently, the only carbapenem included in the guidelines for treating *Mab* infections is imipenem, which has a CLSI breakpoint for susceptibility at <4 μg/ml, intermediate activity at 8-16 μg/ml, and resistance at >32 μg/ml (31). Sulopenem *in vitro* activity was comparable to sulopenem in this study. Moreover, the addition of 4 μg/ml of CXM led to a significant further reduction in MICs in over 90% of the clinical isolates. These data suggest its potential use as an oral step-down therapy following an initial induction phase. Notably, both sulopenem and CXM are available in oral formulations. Similar synergy with combinations of tebipenem and avibactam was observed in recent study (30).

The time-kill analysis provided further support of the MIC outcomes, corroborating the *in vitro* efficacy of sulopenem in killing of *Mab* for both ATCC 19977 and the clinical isolate *Mab*122. Augmentation of the potency of sulopenem was observed by the addition of CXM or DBOs such as AVI or DUR. The synergistic impact of the dual β-lactam combination (sulopenem and CXM) in the SCTK was somewhat diminished compared to that observed in the MIC test. This discrepancy might be attributed to the compensatory action of thermally stable drugs counteracting the effects of thermally unstable drug during the MIC test. Both sulopenem and CXM exhibit limited thermal stability (with CXM being relatively more stable) (30). In the susceptibility testing, we were unable to counterbalance the thermal degradation over the 3-7-day testing period, unlike in the time-kill study where such supplementation was feasible. The practice of supplementing unstable agents to counteract chemical degradation has implications for MIC data interpretation. While the CLSI does not currently stipulate supplementation in susceptibility testing for rapidly growing mycobacteria, pertinent effects on MIC testing in *Mab* have been reported (32).

Macrolides, which historically found common utility in treating *Mab* infections, exhibited notable efficacy against susceptible *Mab* strains, albeit primarily manifesting bacteriostatic effects. In the context of our time-kill assay, a majority of drug regimens, encompassing monotherapies, dual β-lactams, and β-lactam combined with AVI, elicited bacteriostatic effects, with an exception observed upon the addition of DUR, culminating in a bactericidal effect. This combination strategy, yielding a bactericidal impact, holds promise for mitigating elevated relapse rates and suboptimal treatment outcomes. Given that certain antibiotics exhibit limited penetration into bronchial secretions (2), particularly pertinent in *Mab* lung infections, opting for interventions demonstrating bactericidal activities aligns logically.

The assessment of β-lactam efficacy in the SCTK study encompassed an examination involving one wild-type isolate (ATCC 19977) alongside randomly selected two distinct clinical isolates (*Mab* 122 and *Mab* 686). Paralleled patterns of bacterial killing efficacy were discerned in both ATCC 19977 and *Mab* 122. In contrast, growth inhibition was the primary outcome exhibited by *Mab* 686, yielding much lower bacterial killing. Consistently, high MIC value of this isolate was observed. Furthermore, the addition of BLIs failed to elicit enhancements in bacterial killing. Plausible explanations for this phenomenon encompass potential mutations in peptidoglycan synthesis proteins, including Ldts and PBPs, or a plausible reduction in cell envelope permeability. To gain deeper insights, comprehensive investigations are warranted, involving genome sequencing and subsequent comparative analyses between the ATCC reference strain and these clinically derived isolates.

In conclusion, this study unveils the *in vitro* synergy of sulopenem against *Mab*, shedding light on its potential as a therapeutic agent. Additionally, we have postulated a plausible mechanism underpinning its efficacy and have illustrated its synergistic interplay with CXM. These discoveries augment the ever-expanding roster of β-lactam compounds demonstrating remarkable effectiveness against *Mab*. Given the growing prevalence and therapeutic challenges posed by NTM infections, particularly *Mab*, the imperative for novel drug development looms large, as healthcare professionals worldwide grapple with the dearth of definitive treatment protocols. Remarkably, most of the synergistic β-lactams scrutinized herein boast well-established pharmacokinetic/pharmacodynamics profiles and a decades-long safety record. Thus, initiating an adaptive clinical trial to scrutinize the comparative efficacy of dual β-lactam therapy against the current standard of care assumes paramount significance.

## Materials and Methods

### Bacterial strains, Antibiotics, and reagents

The 54 clinical isolates analyzed in this study were obtained from de-identified patients. To ensure consistency and reliability, we specifically selected isolates from our well-characterized whole genome-sequenced clinical isolates belonging to the *Mycobacterium abscessus* subspecies *abscessus*. Of these, 34 isolates were sourced from National Jewish Health, while 11 and 9 isolates were obtained from University Hospitals Cleveland Medical Center and Cleveland Metrohealth, respectively. Additionally, ATCC 19977 was acquired from the American Type Culture Collection (ATCC). The active ingredient CXM (CXM) salts and avibactam (AVI) were purchased from AchemBlock, sulopenem was sourced from Iterum Therapeutics, and durlobactam (DUR) was provided from Entasis Therapeutics. All β-lactams and BLIs were prepared in sterile, distilled water or in broth and filter-sterilized with a 0.22 μm PES syringe filter.

### *In vitro* Susceptibility Testing and Combination Studies. Minimum Inhibitory Concentration (MIC) Determination

*In vitro* susceptibility testing. Minimum inhibitory concentrations (MICs) of sulopenem, CXM was determined using microdilution. Approximately 5 x 105 colony-forming units (CFU) per milliliter were inoculated into Middlebrook 7H9 broth supplemented with 10% (vol/vol) oleic albumin dextrose catalase (OADC) and 0.05% (vol/vol) Tween 80. When CXM is combined with sulopenem, the CXM was added at fixed concentration of 4 μg/mL to serial dilutions of sulopenem. Isolates were incubated with test agents at 30°C for 3-7 days, and MIC was defined as lowest antibiotic concentration that prevented visible bacterial grow. We use active moiety compounds, i.e cefuroxime salt and not cefuroxime axetil.

### Cloning and Purification of LDTs, DDC, PBP B, and Bla_Mab_

Cloning and purification of Ldts (Ldt_Mab2_ – Ldt_Mab4_), DDC, PBP B, and Bla_Mab_ were performed as previously described (7). Briefly, truncated sequences of Ldts, DDC, and PBP B (Δ1–41) were generated by Celtek Biosciences and cloned into the pET28(a)+ vector with a TEV (tobacco etch virus) protease cleavage site prior to the start codon of the target protein sequences. Clones were transformed into E. coli BL2 (DE3) and grown to reach 0.6-0.8 at an OD_600_, and protein expression was induced with 0.25 mM isopropyl β-d-1-thiogalactopyranoside (IPTG). After incubation for 18 h at 18°C, cells were harvested and stored at −20°C overnight. Cell pellets were resuspended in buffer containing 50 mM Tris (pH 8.0), 400 mM sodium chloride, and 1 mM Tris (2-carboxyethyl) phosphine hydrochloride (TCEP), followed by sonication and centrifugation. The supernatant was passed through a His Prep FF 16/10 column (GE Healthcare) and washed with 5 column volumes of buffer, and bound protein was eluted with 500 mM imidazole. Eluted protein was subjected to dialysis overnight at 4°C in buffer containing 50 mM Tris (pH 8.0), 150 mM sodium chloride, and 0.5 mM TCEP in the presence of His-tagged TEV protease (ratio of TEV protease to protein was 1:3). To remove the His tag, uncleaved fusion protein, and His-tagged TEV protease, passage over the His Prep FF 16/10 column was performed. Fractions containing Ldt_Mab2_ was pooled, concentrated, and stored in 20% glycerol at −20°C.

### Static drug-concentration time-kill assay

The SCTK studies were conducted over a period of 10 days in Middlebrook 7H9 broth, enriched with 10% (vol/vol) OADC, 0.2% (vol/vol) glycerol, and 0.05% (vol/vol) Tween 80 in duplicate. The efficacy of β-lactams and BLIs was evaluated using ATCC 19977 and two clinical isolates obtained from National Jewish Health (Mab122 and Mab686). The concentrations of β-lactams and BLIs were carefully chosen, considering MIC and clinically attainable levels in patients based on predicted average unbound steady-state plasma concentrations. To offset thermal degradation, a small supplementary dose of sulopenem and CXM was added daily, guided by stability data on β-lactams provided by our collaborator (data not shown) or previously reported (30). The broth was exchanged with fresh broth containing the appropriate drug concentration every 3 days.

Throughout the SCTK studies, an initial inoculum of 10^5.6^ - 10^6.3^ CFU/mL was used. Viable counts were assessed immediately before dosing (referred to as ‘0 h’) and subsequently at 24-hour intervals for a total of 10 days. To eliminate any carry-over of antibiotics, all samples were thoroughly washed twice with sterile saline and subjected to serial 10-fold dilutions to determine viable counts. The viable counting procedure involved sub-culturing 100 μL of either an undiluted sample or an appropriately diluted sample on Middlebrook 7H10 agar plates supplemented with 1% (vol/vol) OADC, 0.2% glycerol, and 0.05% (vol/vol) Tween 80. This method resulted in a counting limit of 1.0 log_10_ CFU/mL, equivalent to a single colony per agar plate.

### Mass spectrometry analysis of Ldts, D,D-carboxypeptidase, PBP B, and Bla_Mab_

10 µg of Ldt_Mab1-4_, DDC, PBP B, and Bla_Mab_ were incubated at room temperature with sulopenem or cefuroxime at a molar ratio of 1:20 for 5 min, 3 hours and 24 hours in 50 mM Tris-HCl, pH 7.5- and 300-mM sodium chloride for a total reaction volume of 20 μl. Reactions were quenched with 10 μL acetonitrile and added to 1 mL 0.1% formic acid in water. Samples were analyzed using a quadrupole time-of-flight (Q-TOF) Waters Synapt-G2-Si electrospray ionization mass spectrometer (ESI-MS) and Waters Acquity H class ultra-performance liquid chromatography (UPLC) with a BEH C18 1.7 µm column (2.1 x 50 mm). The Synapt G2-Si was calibrated with sodium iodide with a 50-2000 m/z mass range. MassLynx V4.1 was used to deconvolute protein peaks. The tune settings for each sample were as follows: capillary voltage at 3 kV, sampling cone at 35 V, source offset at 35, source temperature of 100 °C, desolvation temperature of 500 °C, cone gas at 100 L/h, desolvation gas at 800 L/h, and 6.0 nebulizer. Mobile phase A was 0.1% formic acid in water. Mobile phase B was 0.1% formic acid in acetonitrile. Mass accuracy for this system is ±5 Da.

#### Molecular modeling, Docking and Analysis

The crystal structure of Ldt_Mab2_ (PDB:5UWV) was used to model the enzyme. The missing loop (D301-D313) was reconstructed using the Ldt_Mt2_ (PDB: 6IYW) structure as a template and SWISS-MODEL homology-modelling server accessible via the ExPASy web server (33). The structure was future minimized and prepared as previously describe (11) using Discovery Studio software (BIOVIA DS Client 2020).

The structural model of Ldt_Mab3_ was similarly generated, using Ldt_Mtb5_ (PDB:4Z7A) as template. The structure was minimized using a Conjugate Gradient method, with a RMS gradient of 0.001 kcal/(mol x Å). Generalized Born with a simple Switching (GBSW) solvation model was used, and long-range electrostatics were treated using a Particle Mesh Ewald method with periodic boundary condition. The SHAKE algorithm was applied.

The intact, acyl and fragmented sulopenem was built and docked into the active site of Ldt_Mab2_ and Ldt_Mab3_ transpeptidases structures. The CDOCKER protocol was used to dock the compounds into the active site of Ldt_Mab2_ and Ldt_Mab3_. The protocol uses a CHARMm-based molecular dynamics (MD) scheme to dock ligands into a receptor binding site as previously described (11). The generated poses were analyzed and the best ranked were used to create the Michaelis-Menten and acyl-enzyme complexes and were further minimized. To equilibrate the structure, a medium-long molecular dynamic simulation (1ns) was performed, using a NAMD protocol (11, 34).

---

**Supplement Fig. 1.**
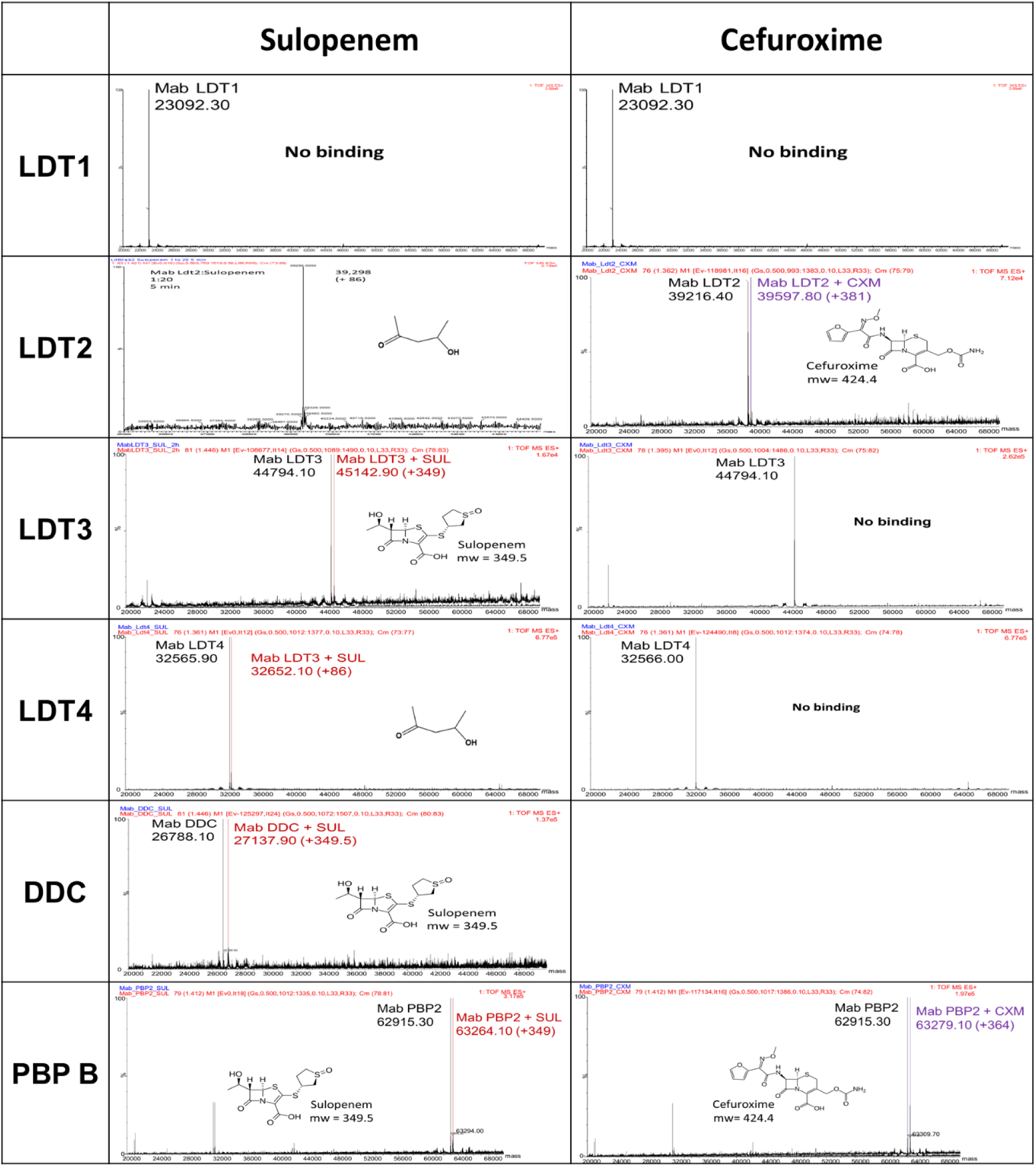
Intact-protein UPLC-IM-MS analysis reveals that sulopenem binds to Ldt_Mab2_, Ldt_Mab4_, DDC, and PBP B within a 5-min. Binding of sulopenem to Ldt_Mab3_ exhibits a binding time exceeding 2 hours. On the other hand, cefuroxime shows binding only towards Ldt_Mab2_ and PBP B. The depicted figure illustrates the interactions, where the black bars represent apo-LDTs, DDC, and PBP B; the red bars indicate the LDTs, DDC, and PBP B-sulopenem adducts, and the purple bars represent the LDTs, DDC, and PBP B-cefuroxime adducts.

**Supplement Fig. 2:**
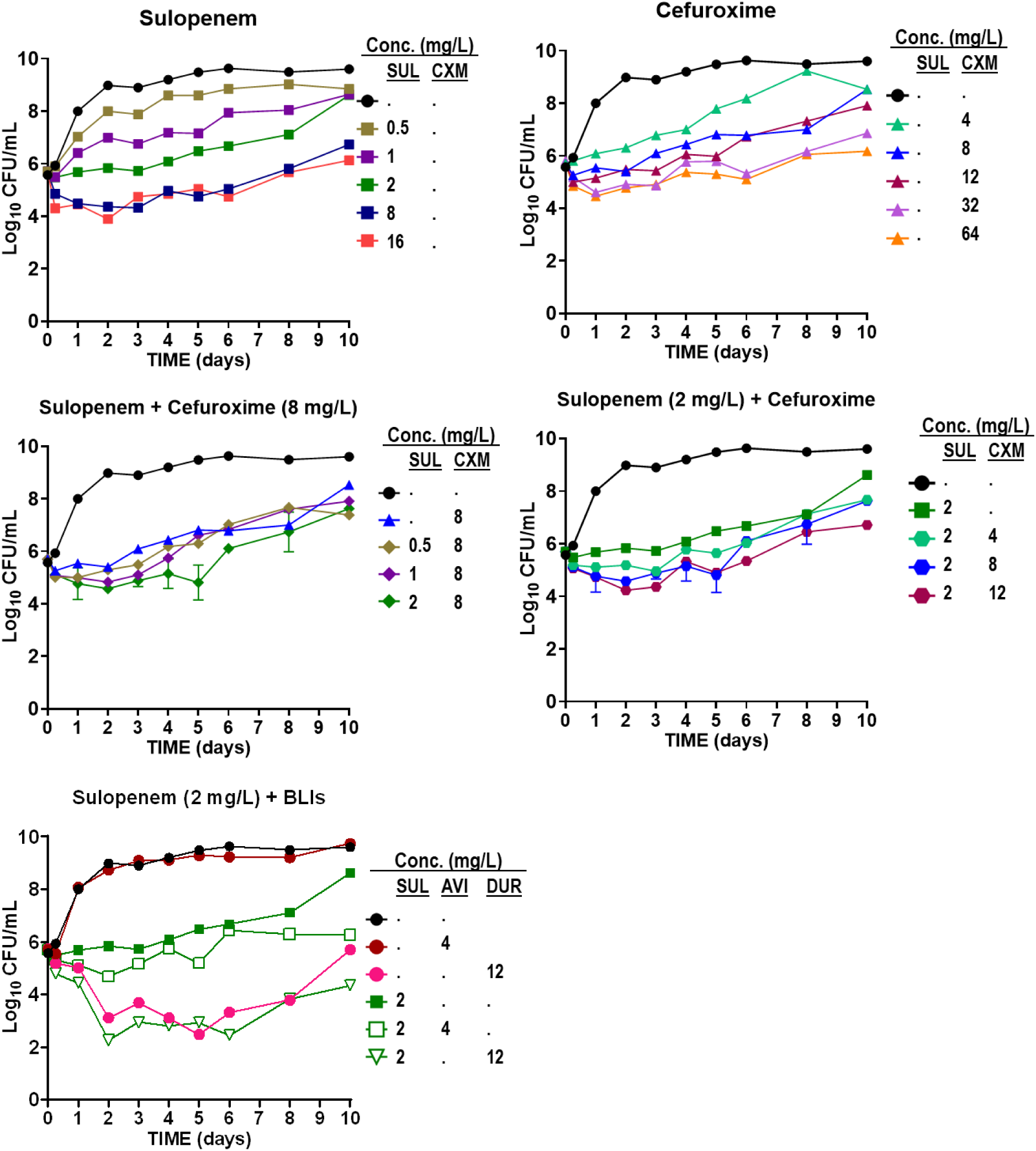
Time-kill curves of sulopenem (A) in monotherapy, cefuroxime (CXM) in monotherapy (B), the combination of sulopenem and CFX (C and D), and sulopenem in the presence of BLIs (avibactam; AVI and durlobactam; DUR) (E) against clinical isolate *Mab* 122.

**Supplement Fig. 3:**
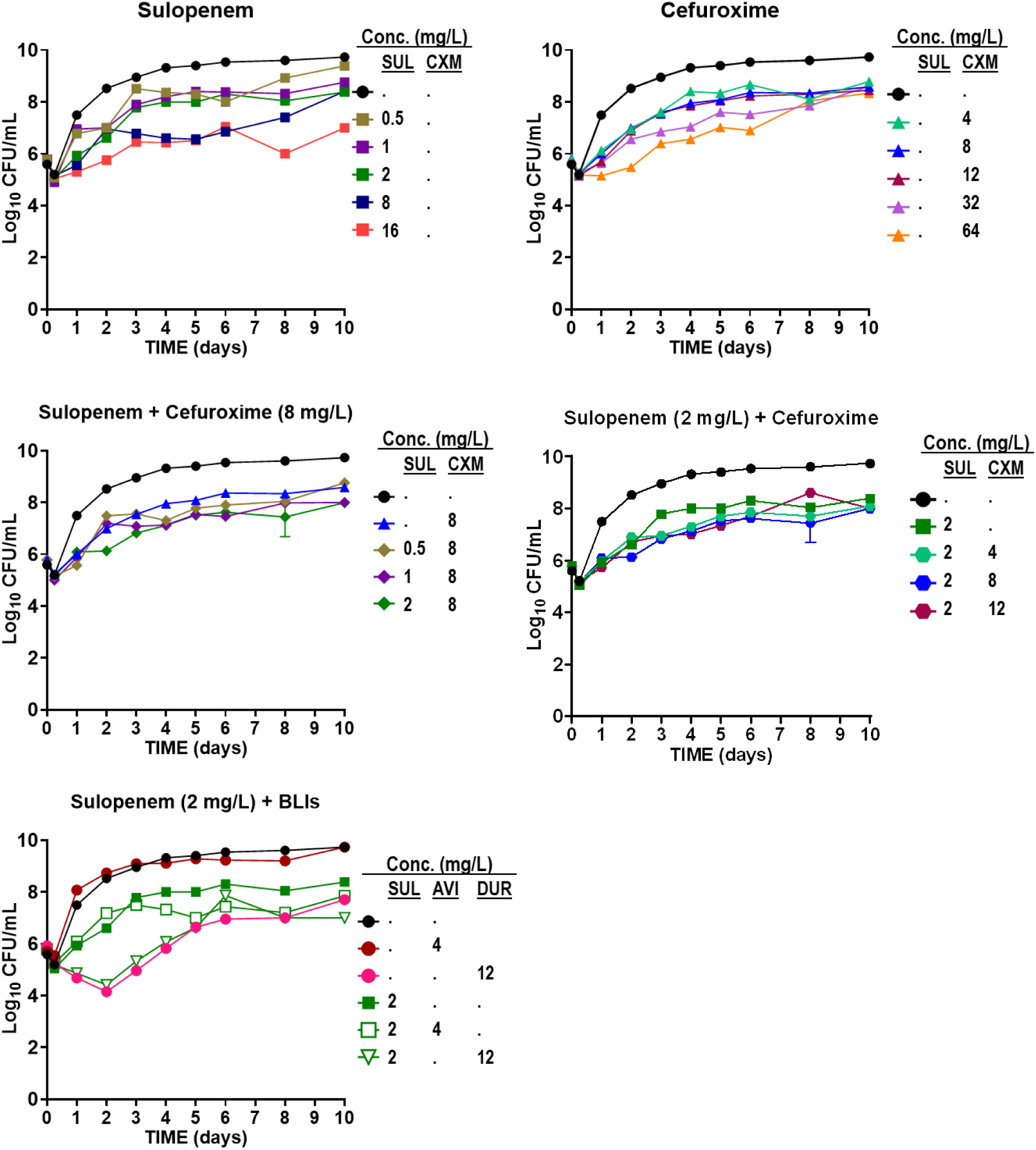
Time-kill curves of sulopenem (A) in monotherapy, cefuroxime (CXM) in monotherapy (B), the combination of sulopenem and CXM (C and D), and sulopenem in the presence of BLIs (avibactam; AVI and durlobactam; DUR) (E) against clinical isolate *Mab* 686.

**Supplement Fig.4:**
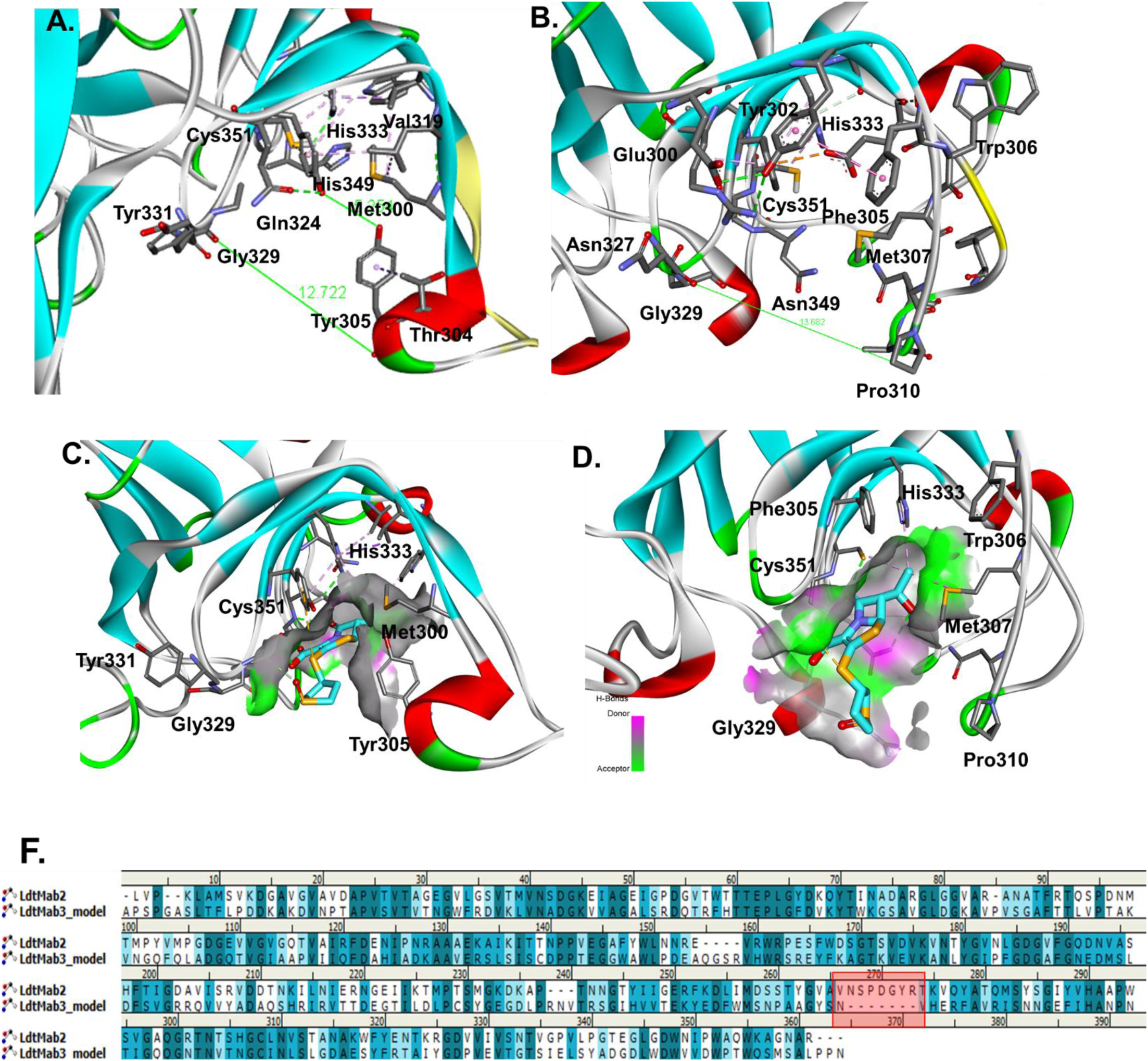
Ldt_Mab2_ (A, C) *vs.* Ldt_Mab3_ (B, D) active site variability and shape. The deletion and insertion into the loops of the active site are colored yellow and red for sequnce alingnment representation (F).

## References

1. Nessar R, Cambau E, Reyrat JM, Murray A, Gicquel B. Mycobacterium abscessus: a new antibiotic nightmare. J Antimicrob Chemother. 2012;67(4):810–8.

2. Johansen MD, Herrmann JL, Kremer L. Non-tuberculous mycobacteria and the rise of Mycobacterium abscessus. Nat Rev Microbiol. 2020;18(7):392–407.

3. Griffith DE, Daley CL. Treatment of Mycobacterium abscessus Pulmonary Disease. Chest. 2022;161(1):64–75.

4. Dousa KM, Kurz SG, Bark CM, Bonomo RA, Furin JJ. Drug-Resistant Tuberculosis: A Glance at Progress and Global Challenges. Infect Dis Clin North Am. 2020;34(4):863–86.

5. Soroka D, Dubee V, Soulier-Escrihuela O, Cuinet G, Hugonnet JE, Gutmann L, et al. Characterization of broad-spectrum Mycobacterium abscessus class A beta-lactamase. J Antimicrob Chemother. 2014;69(3):691–6.

6. Daley CL, Iaccarino JM, Lange C, Cambau E, Wallace RJ, Jr., Andrejak C, et al. Treatment of nontuberculous mycobacterial pulmonary disease: an official ATS/ERS/ESCMID/IDSA clinical practice guideline. Eur Respir J. 2020;56(1).

7. Dousa KM, Nguyen DC, Kurz SG, Taracila MA, Bethel CR, Schinabeck W, et al. Inhibiting Mycobacterium abscessus Cell Wall Synthesis: Using a Novel Diazabicyclooctane beta-Lactamase Inhibitor To Augment beta-Lactam Action. mBio. 2022;13(1):e0352921.

8. Kurepina N, Chen L, Composto K, Rifat D, Nuermberger EL, Kreiswirth BN. CRISPR Inhibition of Essential Peptidoglycan Biosynthesis Genes in Mycobacterium abscessus and Its Impact on beta-Lactam Susceptibility. Antimicrob Agents Chemother. 2022;66(4):e0009322.

9. Akusobi C, Benghomari BS, Zhu J, Wolf ID, Singhvi S, Dulberger CL, et al. Transposon mutagenesis in Mycobacterium abscessus identifies an essential penicillin-binding protein involved in septal peptidoglycan synthesis and antibiotic sensitivity. Elife. 2022;11.

10. Galanis C, Maggioncalda EC, Kumar P, Lamichhane G. Glby, Encoded by MAB_3167c, Is Required for In Vivo Growth of Mycobacteroides abscessus and Exhibits Mild beta-Lactamase Activity. J Bacteriol. 2022;204(5):e0004622.

11. Dousa KM, Kurz SG, Taracila MA, Bonfield T, Bethel CR, Barnes MD, et al. Insights into the l,d-Transpeptidases and d,d-Carboxypeptidase of Mycobacterium abscessus: Ceftaroline, Imipenem, and Novel Diazabicyclooctane Inhibitors. Antimicrob Agents Chemother. 2020;64(8).

12. Kumar P, Chauhan V, Silva JRA, Lameira J, d’Andrea FB, Li SG, et al. Mycobacterium abscessus l,d-Transpeptidases Are Susceptible to Inactivation by Carbapenems and Cephalosporins but Not Penicillins. Antimicrob Agents Chemother. 2017;61(10).

13. Nguyen DC, Dousa KM, Kurz SG, Brown ST, Drusano G, Holland SM, et al. "One-Two Punch": Synergistic ss-Lactam Combinations for Mycobacterium abscessus and Target Redundancy in the Inhibition of Peptidoglycan Synthesis Enzymes. Clin Infect Dis. 2021;73(8):1532–6.

14. Alahmdi B, Dousa KM, Kurz SG, Kaufman A, Bonomo RA, Taimur S. Eradicating Pulmonary Mycobacterium abscessus: The Promise of Dual beta-Lactam Therapy. Open Forum Infect Dis. 2023;10(6):ofad312.

15. Wolf AB, Money KM, Chandnani A, Daley CL, Griffith DE, Chauhan L, et al. Mycobacterium abscessus Meningitis Associated with Stem Cell Treatment During Medical Tourism. Emerg Infect Dis. 2023;29(8):1655–8.

16. Moguillansky N, DeSear K, Dousa KM. A 40-Year-Old Female With Mycobacterium abscessus Successfully Treated With a Dual Beta-Lactam Combination. Cureus. 2023;15(6):e40993.

17. Story-Roller E, Maggioncalda EC, Cohen KA, Lamichhane G. Mycobacterium abscessus and beta-Lactams: Emerging Insights and Potential Opportunities. Front Microbiol. 2018;9:2273.

18. Zhanel GG, Pozdirca M, Golden AR, Lawrence CK, Zelenitsky S, Berry L, et al. Sulopenem: An Intravenous and Oral Penem for the Treatment of Urinary Tract Infections Due to Multidrug-Resistant Bacteria. Drugs. 2022;82(5):533–57.

19. Okamoto K, Gotoh N, Nishino T. Pseudomonas aeruginosa reveals high intrinsic resistance to penem antibiotics: penem resistance mechanisms and their interplay. Antimicrob Agents Chemother. 2001;45(7):1964–71.

20. Kumar P, Kaushik A, Lloyd EP, Li SG, Mattoo R, Ammerman NC, et al. Non-classical transpeptidases yield insight into new antibacterials. Nat Chem Biol. 2017;13(1):54–61.

21. Lohans CT, Chan HTH, Malla TR, Kumar K, Kamps J, McArdle DJB, et al. Non-Hydrolytic beta-Lactam Antibiotic Fragmentation by l,d-Transpeptidases and Serine beta-Lactamase Cysteine Variants. Angew Chem Int Ed Engl. 2019;58(7):1990–4.

22. Gupta R, Al-Kharji N, Alqurafi MA, Nguyen TQ, Chai W, Quan P, et al. Atypically Modified Carbapenem Antibiotics Display Improved Antimycobacterial Activity in the Absence of beta-Lactamase Inhibitors. ACS Infect Dis. 2021;7(8):2425–36.

23. Batchelder HR, Zandi TA, Kaushik A, Naik A, Story-Roller E, Maggioncalda EC, et al. Structure-Activity Relationship of Penem Antibiotic Side Chains Used against Mycobacteria Reveals Highly Active Compounds. ACS Infect Dis. 2022;8(8):1627–36.

24. Brammer Basta LA, Ghosh A, Pan Y, Jakoncic J, Lloyd EP, Townsend CA, et al. Loss of a Functionally and Structurally Distinct ld-Transpeptidase, LdtMt5, Compromises Cell Wall Integrity in Mycobacterium tuberculosis. J Biol Chem. 2015;290(42):25670–85.

25. Lopeman RC, Harrison J, Rathbone DL, Desai M, Lambert PA, Cox JAG. Effect of Amoxicillin in combination with Imipenem-Relebactam against Mycobacterium abscessus. Sci Rep. 2020;10(1):928.

26. Pandey R, Chen L, Manca C, Jenkins S, Glaser L, Vinnard C, et al. Dual beta-Lactam Combinations Highly Active against Mycobacterium abscessus Complex In Vitro. Mbio. 2019;10(1).

27. Dubee V, Bernut A, Cortes M, Lesne T, Dorchene D, Lefebvre AL, et al. beta-Lactamase inhibition by avibactam in Mycobacterium abscessus. J Antimicrob Chemother. 2015;70(4):1051–8.

28. Story-Roller E, Maggioncalda EC, Lamichhane G. Select beta-Lactam Combinations Exhibit Synergy against Mycobacterium abscessus In Vitro. Antimicrob Agents Chemother. 2019;63(4).

29. Khalid M Dousa M, David C Nguyen, MD, Sebastian G Kurz, MD, Magdalena A Taracila, MS, Christopher Bethel, MS, and Robert A Bonomo, MD. 1572. Combination Cefuroxime and Sulopenem is active in vitro against Mycobacterium abscessus. Open Forum Infect Dis. 2020.

30. Negatu DA, Zimmerman MD, Dartois V, Dick T. Strongly Bactericidal All-Oral beta-Lactam Combinations for the Treatment of Mycobacterium abscessus Lung Disease. Antimicrob Agents Chemother. 2022;66(9):e0079022.

31. Froberg G, Maurer FP, Chryssanthou E, Fernstrom L, Benmansour H, Boarbi S, et al. Towards clinical breakpoints for non-tuberculous mycobacteria -Determination of epidemiological cut off values for the Mycobacterium avium complex and Mycobacterium abscessus using broth microdilution. Clin Microbiol Infect. 2023;29(6):758–64.

32. Rominski A, Schulthess B, Muller DM, Keller PM, Sander P. Effect of beta-lactamase production and beta-lactam instability on MIC testing results for Mycobacterium abscessus. J Antimicrob Chemother. 2017;72(11):3070–8.

33. Guex N, Peitsch MC, Schwede T. Automated comparative protein structure modeling with SWISS-MODEL and Swiss-PdbViewer: a historical perspective. Electrophoresis. 2009;30 Suppl 1:S162–73.

34. Phillips JC, Braun R, Wang W, Gumbart J, Tajkhorshid E, Villa E, et al. Scalable molecular dynamics with NAMD. J Comput Chem. 2005;26(16):1781–802.

